# Magnetically stimulated cryogels to enhance osteogenic and chondrogenic differentiation of stem cells

**DOI:** 10.1101/2021.02.15.431106

**Authors:** Sedat Odabas, Atakan Tevlek, Berkay Erenay, Halil Murat Aydın, Aysun Kılıç Süloglu, Atiye Seda Yar Saglam, Bora Garipcan

## Abstract

Cells can respond to the physical stimulus that comes from their micro-environments. There are several strategies to alter cell behavior. Several tissues like bone and cartilage, which are the point of interest of regenerative medicine, are under significant degrees of mechanical stress in real life. Within this stress, the arising mechanotransduction effect may trigger several behavioral responses on cells. As a novel and efficient way, magnetic nanoparticles can be used to make such a mechanotransductive effect on cells.

In this study, pre-functionalized Fe_3_O_4_ superparamagnetic magnetite nanoparticles were synthesized and used to fabricate gelatin-based magnetic cryogels. Cell growth, tissue-specific metabolic activities, differentiation potential to the bone, and cartilage under static magnetic field at different magnetic field strength (1000-4000G) were investigated. Results indicated that there was a better induction in considerable higher magnetic field among all others and magnetic cryogels helps to mediate mesenchymal stem cell behaviour, promote their growth and induce osteogenic and chondrogenic differentiation.

## 1. Introduction

Regenerative medicine focuses on repair, regeneration, or replacement of damaged or lost tissues and organs. In practice, biocompatible and biodegradable three-dimensional scaffolds can be used to help cells (stem cells or other types of tissues own cells) to adhere, proliferate or differentiate when necessary (Hutmacher et al., 2000; O’Brien et al., 2011). Although there are significant advances in regenerative medicine and scaffold fabrication techniques, still bringing cells together with a plain scaffold does often faced the problem of insufficient regeneration or repair (Williams et al., 2004; Ikada et al., 2006).

Growth factors may be a solution to deal with these challenges by inducing cells to proliferate or differentiate. Administration of growth factors can be achieved by direct injection to the side, embedding to the scaffold for controlled release, or by viral or non-viral transfection of the cell to produce cells own factors (Lee et al., 2011; Inci et al., 2014; Vural et al., 2017; Roca et al., 2018). However, the irregular release, insufficient administration or efficacy, hypertrophic growth, immunological response in treatment are still drawbacks of using growth factors (Aravamudhan et al., 2013; Whitney et al., 2017).

The microenvironment of which cells are communicated within biological organisms has a significant role in cell behavior that may affect tissue regeneration. Cells can interact with each other and their microenvironment through transmembrane proteins, biochemical, and biophysical cues. Studies show that mechanical stress on constant/varying degrees of loads can determine the cell fate, affect cell attachment and proliferation patterns and functions of specific tissues such as bone and cartilage. (Lele et al., 2007; Bilgen et al., 2016; Gómez-González, et al., 2020).

Cells and tissues are constantly under the influence of many physical forces such as hydrostatic pressure, shear stress, tension, and compression forces. To mimic physical stimulatory forces, classical or custom-designed bioreactors can be used. Bioreactors can overcome the diffusion limitations and allow mass transport of the nutrients in three dimensions. On the other hand, shear stress is a troubling issue for cells. Cells may be sensitive to any perfusion or recirculation in the system (Rauh et al. 2011; Yeatts et al., 2011; Inga et al., 2011).

For decades, magnetic particles have been used for industrial, therapeutic, or biomedical applications such as separation and purification, electronics, catalysis technology, drug release, as well as molecular recognition and imaging (McBain et al. 2008; Odabas et al., 2008; Cardosa et al., 2017; Baresel et al., 2019; Jin et al., 2021). Generating the physical stress by using a particulate system may also be a well-driven strategy to mediate the cell behavior. One of the important approaches is using magnetic particles sole or combining with a proper scaffold (Li et al., 2019; Goranov et al., 2020). Particle vibration under a certain magnetic field may generate mechanical stress that can detect by mechanosensors in the cell membrane. Eventually, cell may convert this stress into chemical signals that can alter its metabolic activities such as gene expression, protein synthesis, cellular phenotype, and signaling.

Several tissues like bone and cartilage, which are the point of interest of regenerative medicine, are under significant degrees of mechanical stress in real life. Magnetically stimulated scaffolds could be a good candidate for hard tissue engineering. Apart from the recent literature, here in this study, we demonstrate comparative results of osteogenic and chondrogenic differentiation of mesenchymal stem cells on magnetic cryogels under two different magnetic strengths.

## 2. Materials and methods

All materials were purchased from Sigma-Aldrich (Germany) unless otherwise mentioned and used as instructed.

### 2.1. Preparation of magnetic cryogels

Magnetic cryogels were prepared by using pre-functionalized magnetite particles (Fe_3_O_4_). These particles (MNP) were synthesized and functionalized according to a previous report (Odabas et al., 2008). Briefly, Fe^2+^ and Fe^+3^ salts (1:2 M) were co-precipitated with NaOH (3M) under N2 atmosphere at 80°C. Later on, synthesized nanoparticles were coated with a cationic polymeric layer comprising PEG/Styrene/DMAPM (N-[3-(Dimethylamino)-propyl]methacrylamide) via emulsifier free emulsion polymerization (Guven et al., 2004).

Later on, magnetic cryogels were prepared with a slight modification of a previous report (Odabas et al., 2016). Briefly, gelatin (4 %; w/w) was soaked into 0.05 M acetic acid and stirred two hours at room temperature. Magnetic nanoparticles were added (5 %; w/w) to gelatin solution under vigorous mixing. This magnetic gelation solution was kept two hours at room temperature and then poured into plastic casts. Samples were kept first at −20°C for two hours and then transferred to −80°C prior to overnight lyophilization.

Lyophilized samples were cross-linked via a well-known NHS/EDC (25mM NHS/50mM EDC) carbodiimide chemistry under an ethanol environment (Vrana et al., 2007). Samples were then washed several times with Ca^2+^ and Mg^+2^ free Phosphate Buffered Saline (D-PBS at pH 7.4), distilled water, and kept at −80°C prior to overnight lyophilization (Supplementary Figure I).

### 2.2 Characterization of magnetic particles and the cryogels

Size and size distribution and the zeta potential of magnetic nanoparticles were characterized by Zeta Sizer (Malvern NanoZS, UK) and Transmission Electron Microscopy (TEM, Tecnai G2 20 TWIN, FEI Company, USA). TEM analysis was performed at an emission voltage of 200 kV and LaB6 electron emitter was used and images.

Magnetic properties by means of magnetization curves of both magnetic particles and the cryogels were evaluated by using Vibrating Sample Measurement (VSM) (S700X SQUID Magnetometer, Cryogenic Limited, UK). All measurements were performed at room temperature.

Structural characterization of the magnetic particles and the cryogels were evaluated by X-Ray Diffractometer (X-RD, Rigaku D/Max 2200 ULTIMAN, Japan). Samples were scanned in 2 Theta with a scan rate of 5°/min.

The chemical composition of both magnetic particles and the cryogels were evaluated by Fourier-transform infrared (FTIR) spectroscopy (Bruker, USA). The measurements were performed in the frequency interval of 4000–400 cm^−1^ with a resolution of 4 cm^−1^.

Morphological properties of the cryogels were evaluated by Scanning Electron Microscope (SEM-FEI, UK). Samples were coated with gold by sputter coating before observation. Overall porosity of the cryogels was calculated by the average 3D porosity values obtained by Micro-CT scanner (Bruker, Skyscan 1272, CTAn software). ImageJ software was used to calculate porosity of the cryogels (Image J version 1.80 NIH, US).

Swelling ratios of the cryogels were determined in designated time points by the following equation where Wd is the dried weight before soaking in PBS and Ws is the swelled weight after soaking in PBS. Degradation of the magnetic cryogels was evaluated according to well-known protocol at DMEM solution containing 0.005% Na-azide under a mild shaking environment at 37°C (Grover et al., 2012).

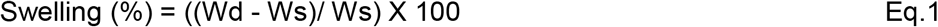

The structural integrity of these cryogels is important for further applications. Therefore magnetic nanoparticles release were monitored under pre-defined condition. Briefly, magnetic cryogels first soaked in 2 ml distilled water. The free release of the magnetic nanoparticles from the cryogels was determined spectroscopically both in dynamic condition (treated under the ultrasonic bath and calculated at 10 – 120 min.) and in static condition (rest in room temperature and calculated up to 5 days).

### 2.3. *In vitro* studies

To assess the possible effect of the magnetic cryogels on mesenchymal stem cell behavior, all experiments were set up under 6 different groups, defined in Table 1.

**Table 1:** Experiment groups for *in vitro* studies

|  | Group ID |
| --- | --- |
| 1 | Plain gelatin cryogels (Gel) |
| 2 | Magnetic gelatin cryogels (M-Gel) |
| 3 | Gel under batch magnet with low magnetic strength (1000Gauss), |
| 4 | M-Gel under batch magnet with low magnetic strength (1000Gauss) |
| 5 | Gel under batch magnet with high magnetic strength (4000Gauss) |
| 6 | M-Gel under batch magnet with high magnetic strength (4000Gauss) |

**Table 2.** Particle release from M-Gel cryogels in static and dynamic conditions over time.

| Dynamic condition |  |  | Static condition |  |  |
| --- | --- | --- | --- | --- | --- |
| Time |  | Weight Loss (%) | Time |  | Weight Loss (%) |
|  | 10 min | 0.80 |  | 1 day | 2.70 |
|  | 30 min | 2.12 |  | 2 days | 2.85 |
|  | 60 min | 3.51 |  | 3 days | 3.07 |
|  | 90 min. | 3.58 |  | 4 days | 3.36 |
|  | 120 min. | 4.02 |  | 5 days | 3.73 |

Cells were cultured in DMEM/F12 supplemented with 10% Fetal Bovine Serum (FBS), 1% L-Glutamine and 1% antibiotic-antimycotic solution named as “basic culture medium”. In differentiation studies, cells were cultured in alpha-MEM supplemented with 10% FBS, 1% antibiotic-antimycotic solution, 10Mm beta-Glycerol Phosphate, 10nM dexamethasone and 100uM ascorbic acid named as “osteogenic medium and were cultured in DMEM-High Glucose supplemented with 10% FBS, 1% antibiotic-antimycotic solution, 10^−7^M dexamethasone, 50uM ascorbic acid, 10ng/ml TGF-β1 and %1 ITS chondrogenic medium.

#### 2.3.1. Cell viability and proliferation

The effect of the magnetic cryogels on cell growth and proliferation were assessed by basic MTT assay. Here, 1×10^5^ cells were seeded on cryogels and incubated in “basic culture medium” for up to 28days. Cell proliferations were evaluated in different groups that mentioned above and also with respect to the Tissue Culture Polystyrene Plate (TCPS) as control.

#### 2.3.2. MSC-Cryogel Interactions

Cell-cryogel interactions and basic cellular morphologies were observed by SEM. Here, 1×10^5^ cells were seeded on cryogels and incubated in “basic culture medium” up to 7days. Samples were then washed several times with D-PBS and then fixed with 2.5% glutaraldehyde for about 30minutes. Samples were dehydrated through an ethanol series and coated with gold prior to observation.

#### 2.3.3. Osteogenic and Chondrogenic Differentiation

With their magnetic ability, magnetic cryogels may induce and improve stem cells ‘differentiation potential through triggering osteogenic and chondrogenic paths. To investigate this phenomena, 1×10^6^ cells were seeded on per cryogels and were cultured in osteogenic and chondrogenic media mentioned above. Osteogenic ad chondrogenic differentiation were evaluated up to 28 days by “Quantitative Analysis” such as tissue-specific histological stainings and gene expression.

In histological staining, samples were stained with Alizarin Red in terms of monitoring osteogenic differentiation and stained with Toluidine Blue for chondrogenic differentiation. Quantitative measurements were performed by the following protocol using cetylpyridinium chloride (CPC) (Maccari et al., 2002; Harvard Medical School, Center for Skeletal Research, 2017). Cell-free cryogels were used for omitting false staining.

Cells were collected for up to 28 days for quantitative gene expression analysis. Total RNA was extracted from the cells using TriReagent (peqGOLD TriFastTM, peqlab, Erlangen, Germany). Total RNA of 1 μg from each sample was reverse transcribed with random hexamers using Transcriptor First Strand cDNA Synthesis Kit (Roche Diagnostics, Germany) according to the manufacturer’s protocol. Quantitative Real-Time polymerase chain reaction (qRT-PCR; Light-Cycler 480 instrument Roche Diagnostics, Mannheim, Germany) analysis was performed for Collagen Type I (COL1A1), Alkaline Phosphatase (ALP), Osteonectin (OSN) gene expression for osteogenic differentiation and for Collagen Type I (COL1A1), Collagen Type II (COL2A1), Aggrecan (Acan) and Early Chondrogenesis Factor (SOX-9) gene expressions for chondrogenic differentiation. Glyceraldehyde-3-phosphate dehydrogenase (GAPDH) was used as a control gene and the results were normalized and evaluated using REST (Relative Expression Software Tool) software.

##### Statistical analysis

The changes in mRNA levels were compared by Relative Expression Software Tool (REST^®^ 2009 v2.013) using 2-ΔΔCT relative quantitative analysis and also confirmed by GeneGlobe Data Analysis Center (Qiagen). Student t test has been used throughout the study. P-values under 0.05 were considered to show a meaningful statistical significant difference. Each analysis was performed in triplicate.

## 3. Results and Discussion

Superparamagnetic particles have the ability to respond to an external magnetic field. With this ability, they can be used for several biomedical and bioengineering applications. Dobson used them to remote control cell signaling under an external magnetic field (Dobson 2008). Subramanian et al. reported the use of magnetic nanoparticles under gradient magnetic field to remote manipulation of neuroblastoma (SH-SY5Y) and hepatocellular carcinoma (HepG2) cells (Subramanian et al., 2019). Apart from *in vitro* studies, there are also magnetic nanoparticle based *in vivo* trials, especially for drug delivery or cancer treatment purpose, using them to guide and target to local tissues under a certain magnetic field (Sensenig et al., 2012; Malik et al., 2017) However, magnetic scaffolds are more versatile and easy to implement approach to mediate the cells both in vitro and in vivo applications. These scaffolds can either be the synthetic or natural origin and allow to improve cell growth as well as tissue regeneration (Bock et al., 2010 Li et al., 2019; Goranov et al., 2020).

### 3.1 Magnetic particle synthesis and characterization

The co-precipitation of iron salts in a strong base condition is a well-known technique. The magnetic particles synthesized here have an average size of 10 nm and approximately unidimensional (Figure 1A). According to size and size distribution measurement by Dynamic Light Scattering (DLS), the average size reaches 200nm (189,85 ±15,25). DLS basically measures the hydrodynamic radius of a cluster, not a single particle. Here, results were in smaller size values (117,4 ± 8,01) after coating. This is due to the hydrophobic interaction of both magnetite and the methacrylate end groups in the polymer, and coating allows magnetite nuclei into smaller aggregates by separating them. The PDI values of the MNP’s were 0,303 ± 0,085 and became more homogeneous after coating (0,077 ± 0,065). The zeta potential (by means of average surface charge) of sole magnetic particles is almost −12mV (−12,2 ±0,49) and it reached almost + 40mV after coating (+37,8 ± 8,3), which we believe is sufficient enough for better interaction with the cells.

**Figure 1.**
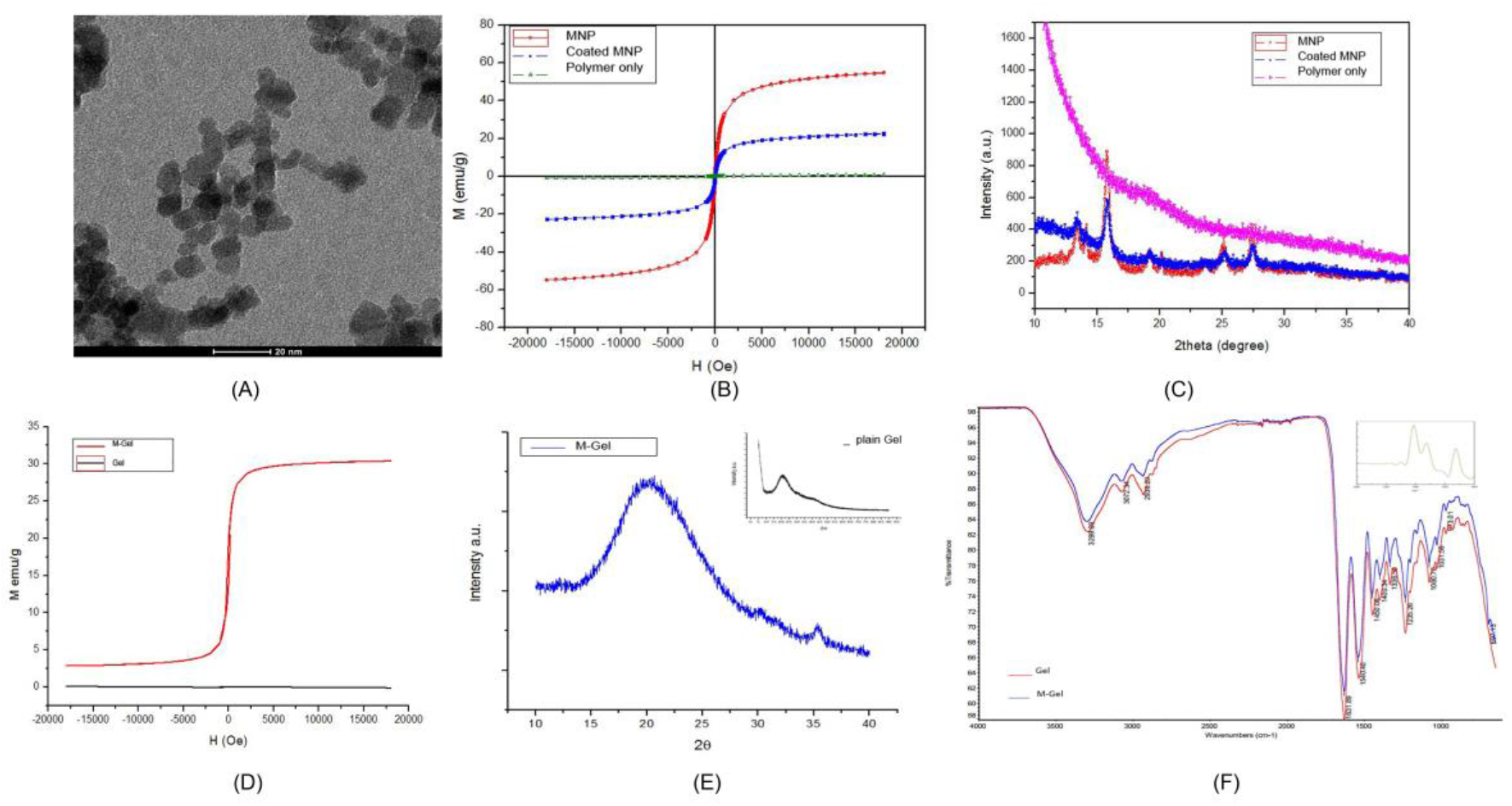
(A) TEM image of MNP (B) VSM analysis of sole MNP and coated MNP (C) X-RD analysis of both sole MNP and coated MNP (D) VSM analysis of the Gel and M-Gel cryogels (E) X-RD analysis of the Gel and M-Gel cryogels (F) FTIR analysis of MNP and cryogels

VSM analysis performed here shows the magnetic properties of the particles in terms of magnetic susceptibility (Hysteresis - H-Oe). According to Figure 1B, the particles showed the superparamagnetic property and had a magnetic saturation of 54.94 emu/g, while commercial samples (Sigma-Aldrich, 637106) showed a max. 65.37 emu/g in a magnetic field of 18000 Oe. The commercial particles have an average size of 50-100 nm, according to the technical sheet. It is expected that the smaller the core size is, the less magnetic strength has. Coating the magnetic particles with a functional layer causes a less magnetic saturation, 22.73 emu/g, at the same magnetic field strength (Figure 1B). This is an expected result as the coating adds extra distance between the magnetic core and the magnetic field applied. However, these functionalized particles still have sufficient magnetic strength to use in cell interactions.

X-RD results depicted in Figure 1C prove the octahedral structure of Fe_3_O_4_ particles. According to the spectrum, the polymer coating does not disturb the crystalline structure of the particles in the comparative analysis of the magnetic nanoparticle, polymer-coated magnetic nanoparticle, and the polymer itself. Besides, the polymer, which does not show crystallinity, expressed itself as a broad and diffuse wave in the analysis.

### 3.2 Magnetic cryogels preparation and characterization

This study is based on the idea of using magnetic cryogels that have the ability to generate physical stress on cells to mediate cell behavior. Moreover, these gels will give a proper response to any magnetic field applied. The comparative VSM result of magnetic moment values per unit mass for Gel and M-Gel cryogels were given in Figure 1D. M-Gel cryogels showed a magnetic saturation of 13.81 emu / g, where Gel cryogel showed zero magnetic saturation (−0,104 emu / g). This result belongs to the block mass of the cryogel rather than the functional magnetic nanoparticle alone. Still, these cryogels in their final form can respond to the magnetic field.

Gelatin in cryogels forms broad flat peaks in X-RD spectrum. Polymeric components of functional magnetic nanoparticles incorporated into the structure express themselves in Figure 1E at 18-20° degrees like gelatin (Mishra et al., 2010; Khan et al., 2015). The software confirmed that the difference between two X-RD reveals the contribution of magnetic nanoparticles.

The comparative spectrum results of the chemical structure characterization performed by FTIR spectroscopy in Gel and M-Gel cryogels were depicted in Figure 1F. According to the spectra, M-Gel findings closely resembled the FTIR spectrum of the Gel cryogels. The benzene ring originating from styrene, which is the main component in the functional polymeric coatings of the magnetic nanoparticles in its content, reveals itself around 698 cm^−1^. In the small inline spectrum, the characteristic peaks of the Fe-O bond structure can be seen. The octahedral and tetrahedral peaks that should be seen in the structure between 630-550 cm^−1^ are around 530 cm^−1^ in the sample. This indicates a small percentage of elemental Fe impurities in the structure partially not goes into reaction. In addition, the characteristic peak of the magnetite octahedral structure with a small amount of elemental Fe impurity is observed around 440 cm^−1^ (Gawali et al. 2021). When the FTIR spectra of MNPs were examined, it was observed that the sample compared with the commercial sample had large spectral compatibility. Spread around 3300cm^−1^ in the spectrum of both samples reflects the humidity-induced -OH stress, peaks around 2200cm^−1^, C-C stresses equivalent to residual organic residues from the environment during washing and drying steps, and small peaks around 1400cm^−1^ reflect C-O stresses caused by atmospheric CO_2_.

In order to compare the effect of cross-linking to the thickness of the samples, cryogel thickness was measured before and after the cross-linking using a standard micrometer. The average thickness of Gel and M-Gel cryogels after cross-linking were 3928 ± 110 μm and 4123 ± 173 μm respectively and that had been 4302 ± 166 μm and 4336 ± 105 μm before. Note that, no significant difference between Gel and M-Gel cryogels in terms of thickness values. EDC / NHS cross-linking is based on the formation of an amide bond between carboxyl and amino groups. Therefore it can be said that gelatin fibers take a tighter structure with the bonds formed during cross-linking (Vrana et. al., 2007).

According to the swelling behavior of cryogels, Gel and M-Gel cryogels had a swelling degree of 563.63 ± 51.92% and 657.31 ± 41.42%, respectively (the graph is not shown). Although the difference between Gel and M-Gel is small, it can be said that magnetic gelatin cryogels retain water better than plain gelatin cryogels. This difference is thought to be due to the hydrophilic structure (PEG-MEM and DMAMP) of the fine functional terpolymer around the magnetic nanoparticles. In addition to swelling properties, biodegradation studies performed on M-Gel cryogels for 28 days are given in Figure 2E, which shows the M-Gel cryogels degraded approximately 40% after one month (Grover et al., 2012). The rate of degradation was fast in the early days and tended to slow down over time. In such biomaterials, the biodegradation regime also depends on the substance concentration, crosslinker concentration, and cross-linking degree (Davison et al., 2014).

**Figure 2.**
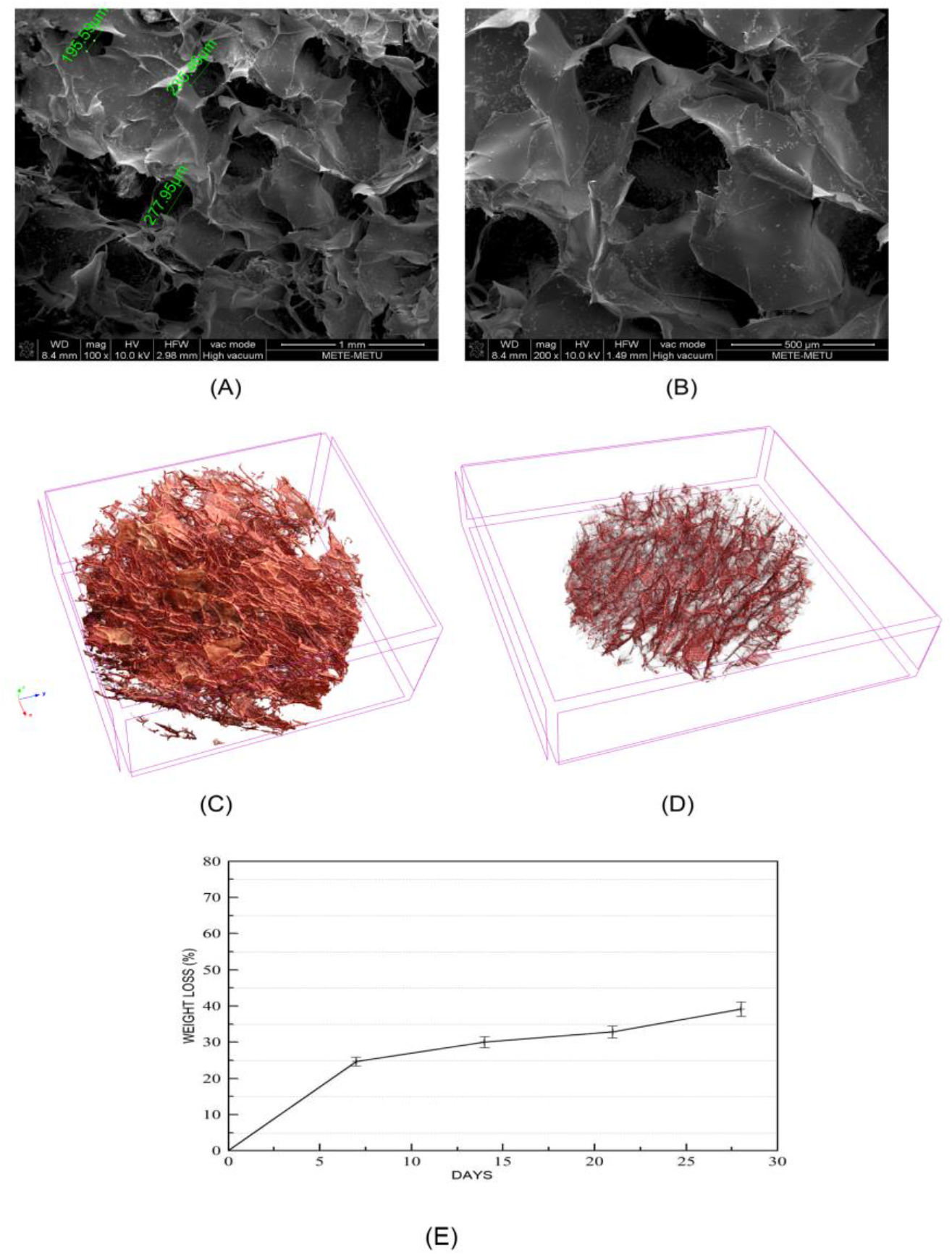
(A-B) SEM, (C-D) Micro-CT image analysis and, (E) biodegradation profile of Gel and M-Gel cryogels.

Particle release test data, performed under static and dynamic conditions for a possible release of magnetic nanoparticles from M-Gel cryogels were depicted in table 4. Particle loss in time is less than 4% (w) in samples both on both minute basis dynamic condition and day basis static condition. It can be speculated that release here is superficial. The M-Gel cryogels here were quite robust and stable in terms of particle-gel interaction, considering the static conditions were closely similar to the real *in vitro* culture conditions.

The porosity and the interconnectivity are crucial for cell attachment and growth (O’Brien et al., 2007). Cryogelation technique is used for the preparation of Gel and M-Gel cryogels. The cryogelation technique is based on the removal of frozen ice crystals from the bulk polymer while forming a pore in the structure. An anisotropic pore structure and porosity emerges in this technique. SEM examination depicted in Figure 2A–2B shows that the cryogels have proper pores and pore sizes of 50-300 um. Microstructural characterization of the cryogels were evaluated by microscope based on micro-computed tomography. Micro-CT was used to visualize the cryogel and reconstructed images were rendered using the CTAn software. It is clearly visible from the representative images depicted in Figure 2C–2D, both Gel and M-Gel cryogels were highly porous. The average porosity of the Gel and M-Gel cryogels were calculated as 80.95 ± 2.47 and 87.96 ± 1.52, respectively. The slightly higher porosity in M-Gel cryogels may be associated with the possible removal of the particles during the preparation.

### 3.3. Cell interaction studies

The superparamagnetic particles used in this study were known to be biocompatible in particular concentrations (Odabas et al., 2008). We investigated the proliferation pattern of MSC as well bone and cartilage oriented differentiation up to 28 days. The proliferative patterns of the cells implanted on cryogels at certain time intervals were determined with MTT assay and were depicted in Figure 3A. TCPS was used as a control. As can be seen from the results, although cells in TCPS showed a rapid proliferation initially, it slowed down at the end of a week due to contact inhibition or due to the lack of enough space for cells to proliferate and reached a plateau at the last stage.

**Figure 3.**
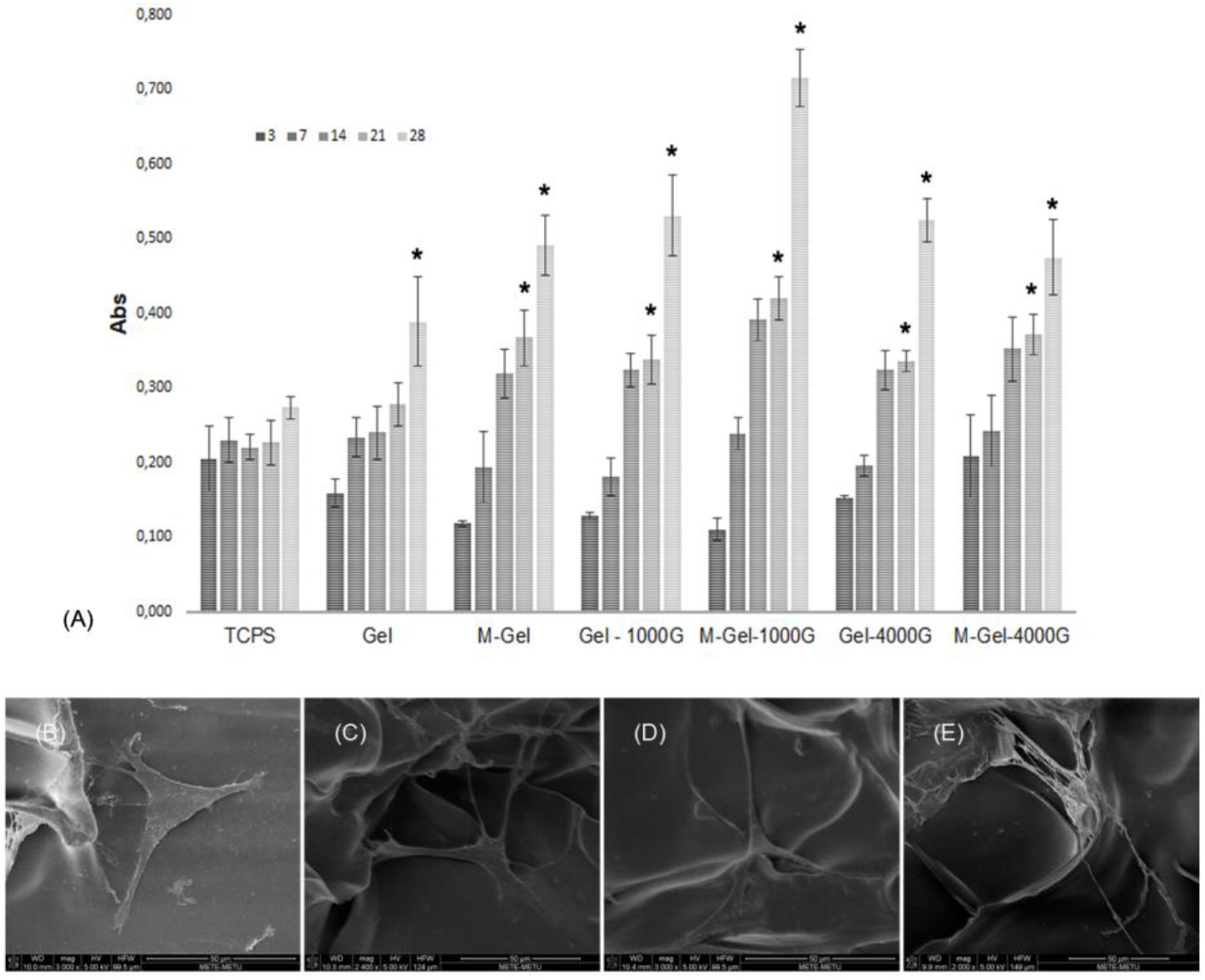
(A) Proliferation pattern of MSCs on cryogels for up to 28days (B-E) SEM images of cells on cryogels at day 7.

On the other hand, the cell densities on cryogel groups increased significantly in the specified time intervals up to 28 days in almost all groups except “Gel” group. There is a rapid increase in M-Gel and batch magnet applied groups, especially for comparably low magnetic strength. In agreement with the previous results, the higher cell densities in batch magnet applied groups may due to the stimulation of the cell division by magneto-mechanical stress generated by vibrated magnetic particles (Zablotskii et al., 2016; Blyakhman et al., 2020).

Cells may behave differently in scaffolds having average pore sizes or scaffold with various gradiated pores regardless of the production technique (Matsiko et al., 2015; Di Luca et al., 2016). Representative SEM images showed (Figure 3B-E) that cells are happy with the cryogels and begin to synthesize their own ECM from the early days of culture. Cells seeded on cryogels were driven into osteogenic and chondrogenic differentiation for up to 28 days. In order to evaluate the differentiation, tissue-specific stainings were performed, and results were depicted in Figure 4.

**Figure 4.**
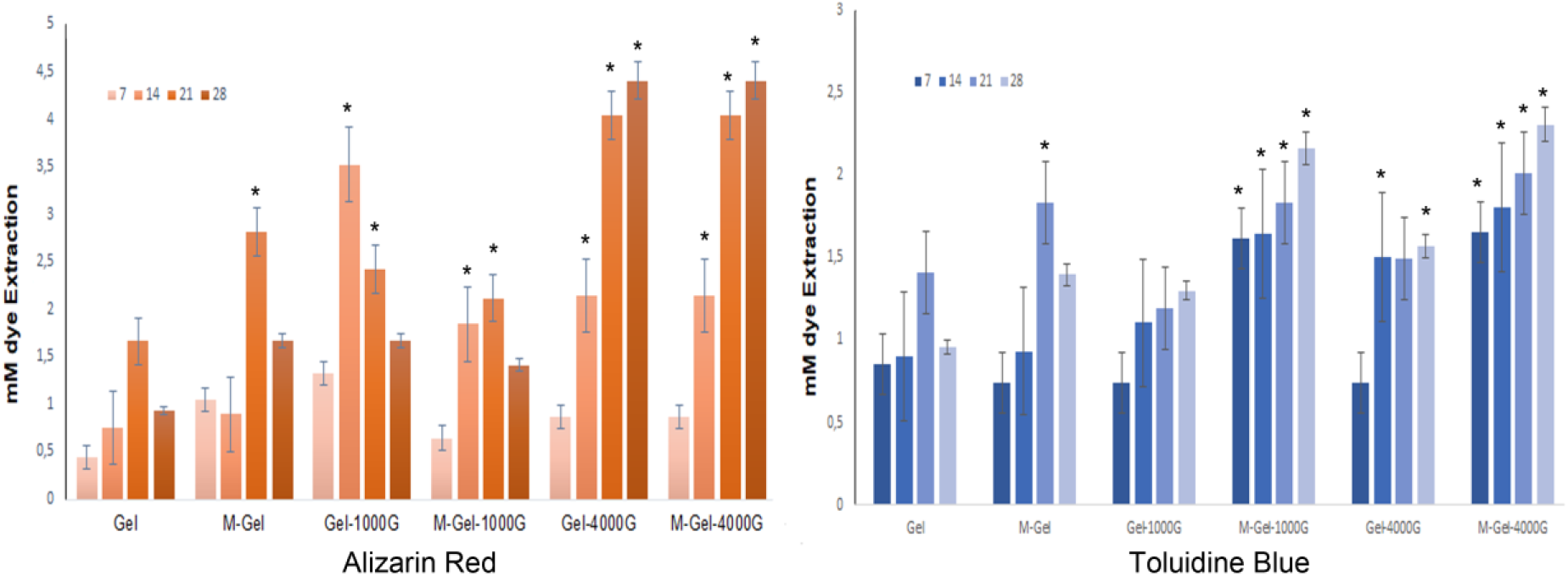
Quantitative analysis of tissue-specific stainings for up to 28days. Alizarin Red staining for osteogenic and Toluidine Blue staining for chondrogenic differentiation.

Alizarin Red staining gives information about the osteogenic behavior of the cells based on calcium deposition. Alizarin Red chelates with calcium ions due to micro-mineralization in the environment and is stained as a red-light orange. The stainings were then quantified by using CPC extraction method and measured spectrophotometrically at 562 nm. Comparing with the plain gelatin cryogels, no significant mineralization was observed on the 7^th^ day, but a significant deposition was observed at batch magnet applied groups after 14^th^ days and significantly increases up to 28^th^ days. The quantitative analysis depicts that there was a better Ca^2+^ deposition in groups under comparably high magnetic strength than other groups.

The chondrogenic differentiation of the cells determined by Quantitative Toluidine Blue staining. Toluidine blue is a glycosaminoglycan fixation dye that gives information about chondro-oriented differentiation. According to the results in Figure 4, there are particular staining intensities in all groups depending on the time interval from day 7 to day 28. Later on the 21^st^ day, almost all groups have similar results. On day 28^th^, compared with plane gelatin cryogels, significant differences were observed in all batch magnet applied groups except Gel-1000G. Similar results with other electromagnetic systems were reported in the literature. For instance, Escobar et al. presented similar GAG findings even with a much higher magnetic field with 2 Tesla (Escobar et al., 2019).

Within the scope of the study, quantitative gene expression analysis was performed to examine the osteogenic and chondrogenic differentiation of mesenchymal stem cells seeded on Gel and M-Gel cryogels in all groups. The aim here is to determine the metabolic activity at the molecular gene level. Initial results were normalized with both GAPDH and the results of day 7. Comparative gene expression study results regarding the differentiation by means of mRNA expression levels are depicted in Figure 5 to Figure 8.

**Figure 5.**
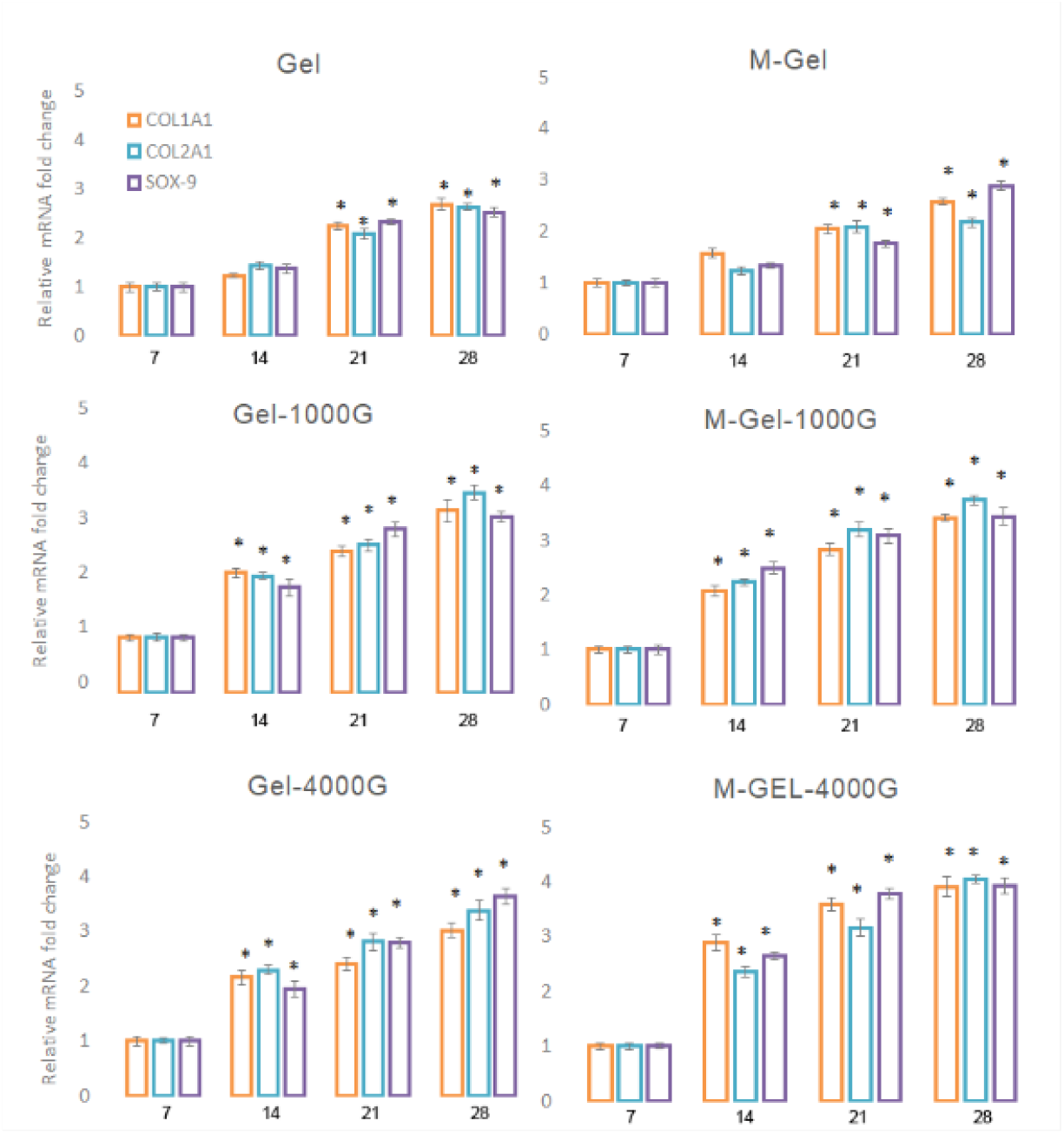
mRNA expression levels of COL1A1, ALP and OSN for osteogenic differentiation. Normalized to GAPDH and to the result of Day 7.

**Figure 6.**
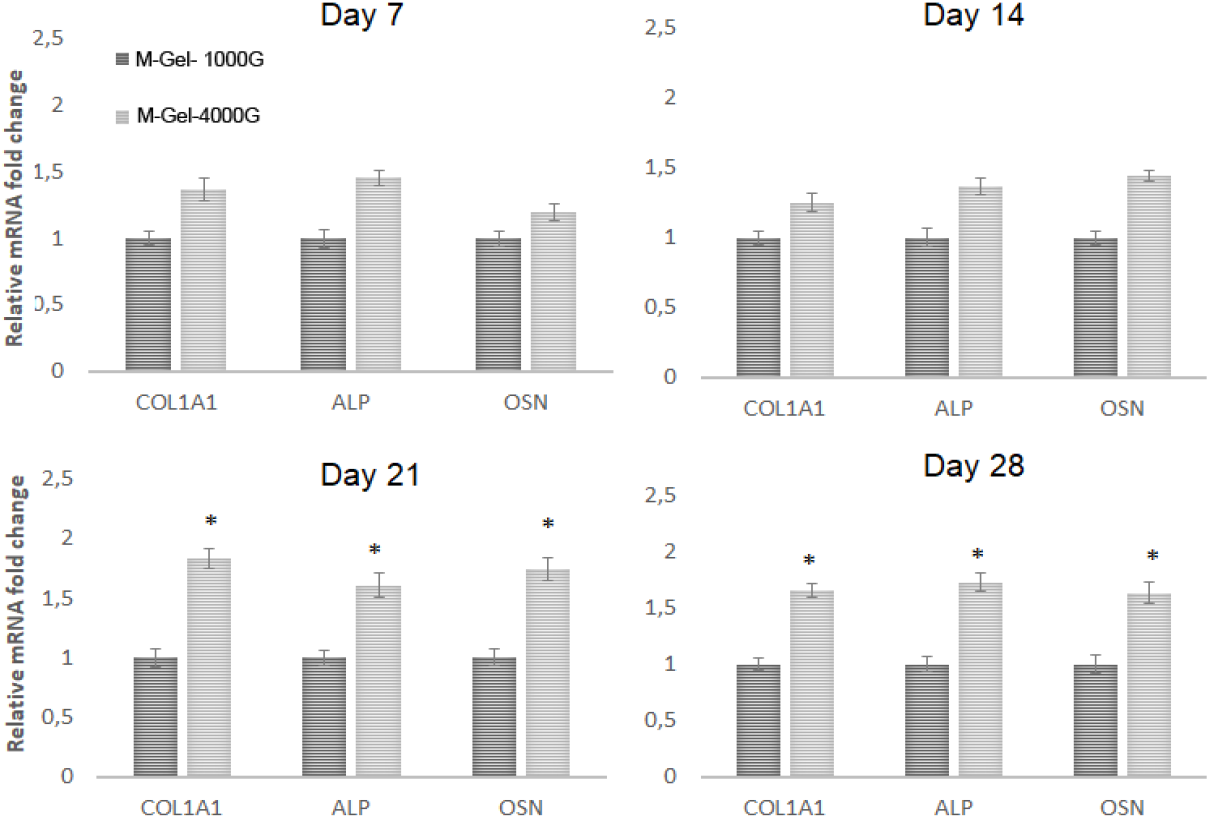
Comparison of magnetic field strength in terms of mRNA expression levels of osteogenic differentiation-related genes of COL1A1, ALP and OSN of cells seeded on magnetic cryogels. Results were normalized to GAPDH.

**Figure 7.**
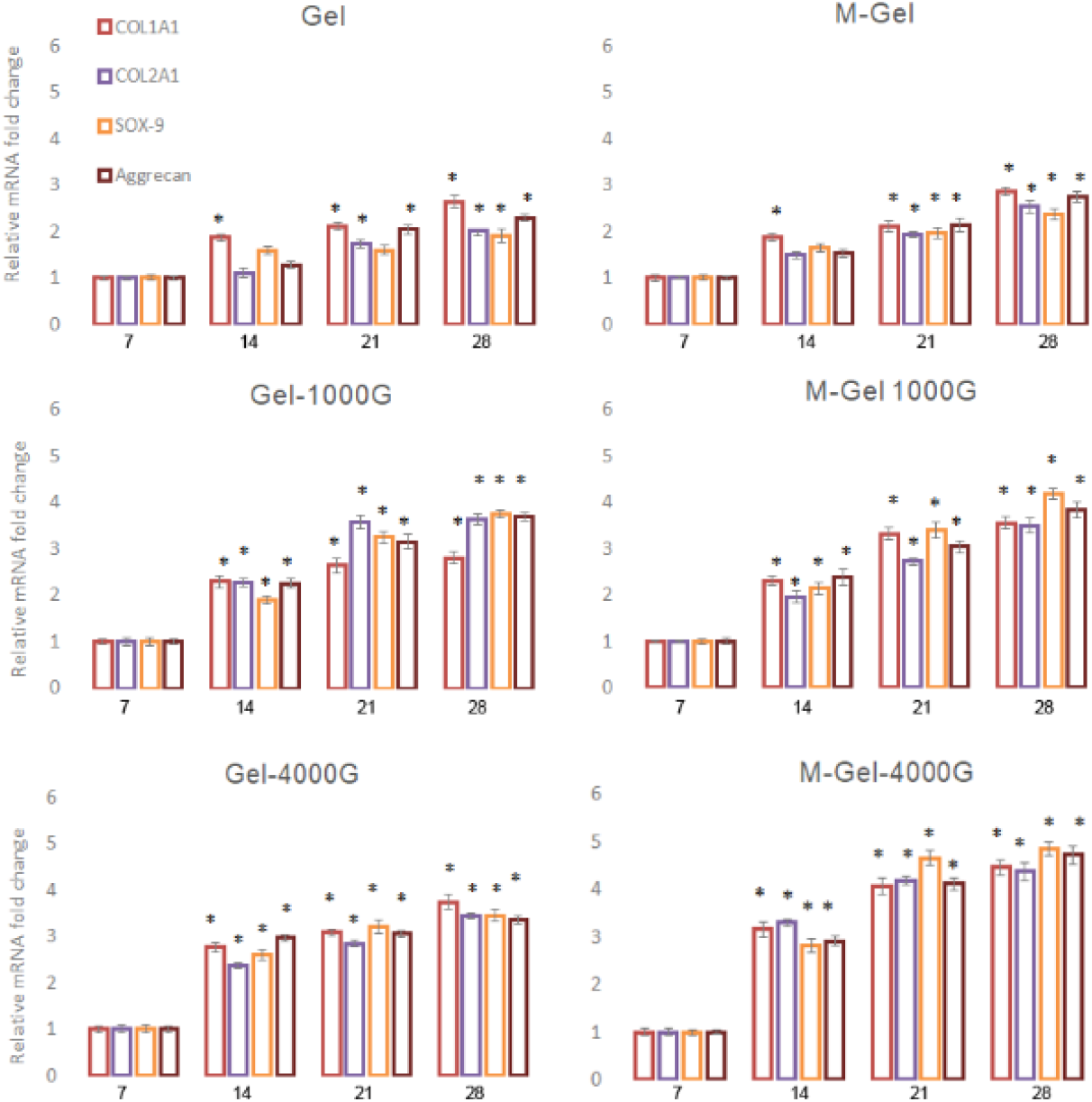
mRNA expression levels of COL1A1, COL1A2, ACAN, SOX-9 for chondrogenic differentiation. Normalized to GAPDH and to the result of Day 7.

**Figure 8.**
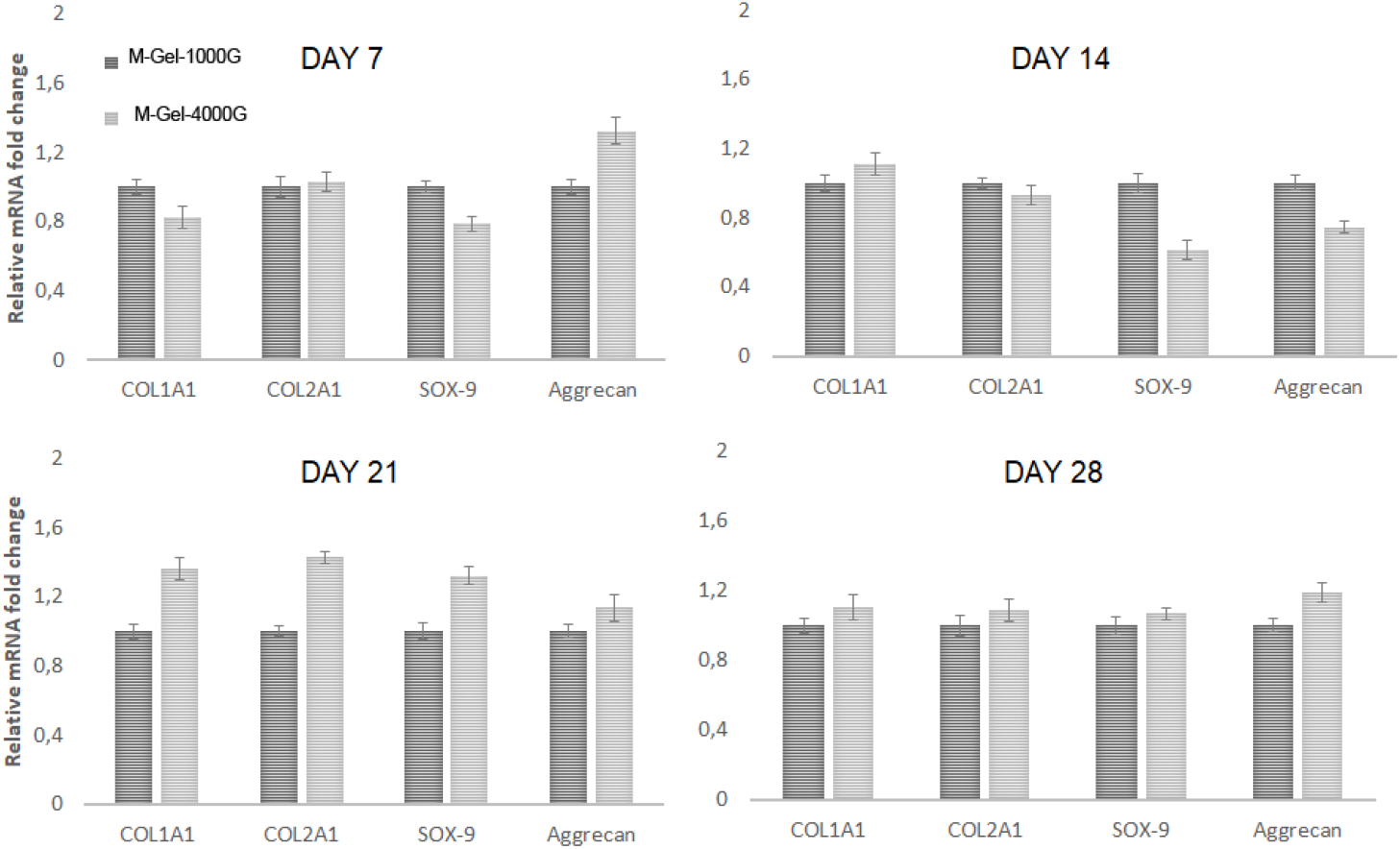
Comparison of magnetic field strength in terms of mRNA expression levels of chondrogenic differentiation-related genes of COL1A1, COL1A2, ACAN, SOX-9 of cells seeded on magnetic cryogels. Results were normalized to GAPDH.

COL1A1, which is related to the synthesis of collagen I, a main component of the ECM, is the main indicator gene for a strong osteogenic differentiation. Alkaline phosphatase (ALP) is an enzyme that is effective in proliferation and mineralization in osteoblasts. Osteonectin (OSN) is a gene related to a glycoprotein secreted from bone cells and playing an important role in early mineralization. Considering the normalized results, there are significant differences in relative mRNA expressions of all batch magnet applied groups. Relative mRNA expressions continue to increase in time for up to 28days for all three genes. The differences are especially high M-Gel-1000G and M-Gel-4000G groups.

As the reports in the literature proposed that, the static magnetic field with magnetic particles could induce osteogenic differentiation (Maredziak et al., 2016, Chang et al., 2020). Here we propose a comparable result of two different magnetic strengths (Figure 6). The results show that there is a difference in the expression of all 3 genes examined in a higher magnetic field (~ 4000 Gauss) group than the lower magnetic field (~ 1000 Gauss). This difference is significant, especially from day 21st, when osteogenic differentiation is most effective in the osteogenic culture medium.

The second part of the gene expression study is the investigation of chondrogenic differentiation of cells seeded on the cryogels. COL1A1 and COL2A1 genes responsible for the synthesis of Collagen Type1 and Type 2, which are the main components of the cartilage ECM. Aggrecan (ACAN) is a cartilage-specific proteoglycan that increases the water retention capacity of the ECM. SOX-9 is an essential transcription factor for early chondrogenesis.

According to the results depicted in Figure 7, there is a significant difference in the relative mRNA expressions of the group for COL1A1 on day 14. Like in osteogenic evaluations, from the day 14^th^ to the day 28^th^, expressions are comparably significant for all four genes in all batch magnet applied group. Here, the expression is much higher in M-Gel-1000G and M-Gel-4000G. Individual evaluation of each group reveals that, all genes expression increased in time and then reached a plateau.

Comparing the magnetic cryogels with the magnetic field applied groups (M-Gel-1000G and M-Gel-4000G) (Figure 8), the results show that the magnetic field does not have a significant effect on differentiation until day 14^th^. Later in the culture, there is a difference in the expression of each four genes examined in the higher magnetic field (~ 4000 Gauss) compared lower magnetic field (~ 1000 Gauss). However, the differences were not found to be significantly in contrary to the osteogenic differentiation results.

## Conclusion

The magnetic field may affect specific cells and alter their metabolic activities. Here, we proposed well-characterized gelatin-based magnetic cryogels and comparative differentiation studies under two different magnetic strength generating by batch neodymium magnets. The findings indicated that cells on magnetic cryogels with the magnetic field applied groups showed better results in proliferation, significantly higher expressions of both osteogenic and chondrogenic directions. However, as a limitation of this study, we can not speculate on the possible effect of the “release from the cryogel-accumulation in the environment” cycle to the cell behavior. Moreover, the dispersion of the magnetic particles in magnetic cryogels is also crucial for generating homogeneous magnetic stress on cells.

Here, the inductive effect to the cells increases *in vitro* with the increase in magnetic strength. As a future perspective, the studies and data conducted here should also be supported by in-vivo animal studies before clinical practice.

## Acknowledgement

This study was funded by the Scientific and Technological Research Council of Turkey (TÜBİTAK) Grant number: 115M522.

## References

Aravamudhan, A., M Ramos, D., Nip, J., Subramanian, A., James, R., D Harmon, M., … & G Kumbar, S. (2013). Osteoinductive small molecules: growth factor alternatives for bone tissue engineering. Current pharmaceutical design, 19(19), 3420–3428.

Atienza-Roca, P., Cui, X., Hooper, G. J., Woodfield, T. B., & Lim, K. S. (2018). Growth factor delivery systems for tissue engineering and regenerative medicine. Cutting-Edge Enabling Technologies for Regenerative Medicine, 245–269.

Baresel, C., Schaller, V., Jonasson, C., Johansson, C., Bordes, R., Chauhan, V., … & Welling, S. (2019). Functionalized magnetic particles for water treatment. Heliyon, 5(8), e02325.

Bilgen, B., & Odabas, S. (2016). Effects of Mechanotransduction on Stem Cell Behavior. Advanced Surfaces for Stem Cell Research, 45–65.

Blyakhman, F. A., Melnikov, G. Y., Makarova, E. B., Fadeyev, F. A., Sedneva-Lugovets, D. V., Shabadrov, P. A., … & Kurlyandskaya, G. V. (2020). Effects of Constant Magnetic Field to the Proliferation Rate of Human Fibroblasts Grown onto Different Substrates: Tissue Culture Polystyrene, Polyacrylamide Hydrogel and Ferrogels γ-Fe2O3 Magnetic Nanoparticles. Nanomaterials, 10(9), 1697.

Bock, N., Riminucci, A., Dionigi, C., Russo, A., Tampieri, A., Landi, E., … & Dediu, V. (2010). A novel route in bone tissue engineering: magnetic biomimetic scaffolds. Acta Biomaterialia, 6(3), 786–796.

Cardoso, V. F., Francesko, A., Ribeiro, C., Bañobre-López, M., Martins, P., & Lanceros-Mendez, S. (2018). Advances in magnetic nanoparticles for biomedical applications. Advanced healthcare materials, 7(5), 1700845.

Chang, C. Y., Lew, W. Z., Feng, S. W., Wu, C. L., Wang, H. H., Hsieh, S. C., & Huang, H. M. (2020). Static magnetic field-enhanced osteogenic differentiation of human umbilical cord-derived mesenchymal stem cells via matrix vesicle secretion. International Journal of Radiation Biology, 96(9), 1207–1217.

Davison, N. L., Barrère-de Groot, F., & Grijpma, D. W. (2014). Degradation of biomaterials. Tissue Engineering, 177–215.

Di Luca, A., Ostrawska, B., Moldero, I.L. et. al. (2016). Gradients in pore size enhance the osteogenic differentiation of human mesenchymal stromal cells in three-dimensional scaffolds, Scientific Reports, 6, 22898.

Dobson, J. (2008). Remote control of cellular behaviour with magnetic nanoparticles. Nature nanotechnology, 3(3), 139–143.

Escobar, J. F., Vaca-González, J. J., Guevara, J. M., Vega, J. F., Hata, Y. A., & Garzón-Alvarado, D. A. (2020). In vitro evaluation of the effect of stimulation with magnetic fields on chondrocytes. Bioelectromagnetics, 41(1), 41–51.

Gawali, S. L., Shelar, S. B., Gupta, J., Barick, K. C., & Hassan, P. A. (2021). Immobilization of protein on Fe_3_O_4_ nanoparticles for magnetic hyperthermia application. International Journal of Biological Macromolecules, 166, 851–860.

Goranov, V., Shelyakova, T., De Santis, R., Haranava, Y., Makhaniok, A., Gloria, A., … & Dediu, V. A. (2020). 3D patterning of cells in magnetic scaffolds for tissue engineering. Scientific reports, 10(1), 1–8.

Grover, C. N., Cameron, R. E., & Best, S. M. (2012). Investigating the morphological, mechanical and degradation properties of scaffolds comprising collagen, gelatin and elastin for use in soft tissue engineering. Journal of the mechanical behavior of biomedical materials, 10, 62–74.

Gómez-González, M., Latorre, E., Arroyo, M., & Trepat, X. (2020). Measuring mechanical stress in living tissues. Nature Reviews Physics, 2(6), 300–317.

Güven, G., Tuncel, A., & Pişkin, E. (2004). Monosized cationic nanoparticles prepared by emulsifer-free emulsion polymerization. Colloid and Polymer Science, 282(7), 708–715.

Harvard Medical School, Center for Skeletal Research, http://www.csrmgh.org/wpcontent/uploads/2017/04/Alizarin_Red-1.pdf. 2017.

Hu, S. H., Liu, T. Y., Tsai, C. H., & Chen, S. Y. (2007). Preparation and characterization of magnetic ferroscaffolds for tissue engineering. Journal of Magnetism and Magnetic Materials, 310(2), 2871–2873.

Hutmacher, D. W. (2000). Scaffolds in tissue engineering bone and cartilage. Biomaterials, 21(24), 2529–2543.

Inci, I., Odabas, S., Vargel, I., Guzel, E., Korkusuz, P., Cavusoglu, T., … & Piskin, E. (2014). Gelatin-hydroxyapatite cryogels with bone morphogenetic protein-2 and transforming growth factor beta-1 for calvarial defects. Journal of Biomaterials and Tissue Engineering, 4(8), 624–631.

Ikada, Y. (2006). Challenges in tissue engineering. Journal of the Royal Society Interface, 3(10), 589–601.

Ingber, D. E. (2002). Mechanical signaling and the cellular response to extracellular matrix in angiogenesis and cardiovascular physiology. Circulation research, 91(10), 877–887.

Jin, Z., Koo, T. M., Kim, M. S., Al-Mahdawi, M., Oogane, M., Ando, Y., & Kim, Y. K. (2021). Highly-sensitive magnetic sensor for detecting magnetic nanoparticles based on magnetic tunnel junctions at a low static field. AIP Advances, 11(1), 015046.

Khan, S., Boateng, J. S., Mitchell, J., & Trivedi, V. (2015). Formulation, characterisation and stabilisation of buccal films for paediatric drug delivery of omeprazole. Aaps Pharmscitech, 16(4), 800–810.

Lee, K., Silva, E. A., & Mooney, D. J. (2011). Growth factor delivery-based tissue engineering: general approaches and a review of recent developments. Journal of the Royal Society Interface, 8(55), 153–170.

Lele, T. P., Sero, J. E., Matthews, B. D., Kumar, S., Xia, S., Montoya-Zavala, M., … & Ingber, D. E. (2007). Tools to study cell mechanics and mechanotransduction. Methods in cell biology, 83, 441–472.

Li, J., Li, Z., Chu, D., Jin, L., & Zhang, X. (2019). Fabrication and biocompatibility of core–shell structured magnetic fibrous scaffold. Journal of biomedical nanotechnology, 15(3), 500–506.

Maccari, F., & Volpi, N. (2002). Glycosaminoglycan blotting on nitrocellulose membranes treated with cetylpyridinium chloride after agarose-gel electrophoretic separation. Electrophoresis, 23(19), 3270–3277.

Malik, A., Tahir Butt, T., Zahid, S., Zahid, F., Waquar, S., Rasool, M., … & Qazi, A. M. (2017). Use of magnetic nanoparticles as targeted therapy: theranostic approach to treat and diagnose cancer. Journal of Nanotechnology, 2017.

Marędziak, M., Śmieszek, A., Tomaszewski, K. A., Lewandowski, D., & Marycz, K. (2016). The effect of low static magnetic field on osteogenic and adipogenic differentiation potential of human adipose stromal/stem cells. Journal of Magnetism and Magnetic Materials, 398, 235–245.

Marijanovic, I., Antunovic, M., Matic, I., Panek, M., & Ivkovic, A. (2016). Bioreactor-based bone tissue engineering. In Advanced Techniques in Bone Regeneration. IntechOpen.

Matsiko, A., Gleeson, J. P., & O’Brien, F. J. (2015). Scaffold mean pore size influences mesenchymal stem cell chondrogenic differentiation and matrix deposition. Tissue Engineering Part A, 21(3-4), 486–497.

McBain, S. C., Yiu, H. H., & Dobson, J. (2008). Magnetic nanoparticles for gene and drug delivery. International journal of nanomedicine, 3(2), 169.

Mishra, R. K., & Ray, A. R. (2011). Synthesis and characterization of poly {N-[3-(dimethylamino) propyl] methacrylamide-co-itaconic acid} hydrogels for drug delivery. Journal of Applied Polymer Science, 119(6), 3199–3206.

O’brien, F. J. (2011). Biomaterials & scaffolds for tissue engineering. Materials today, 14(3), 88–95.

O’Brien, F. J., Harley, B. A., Waller, M. A., Yannas, I. V., Gibson, L. J., & Prendergast, P. J. (2007). The effect of pore size on permeability and cell attachment in collagen scaffolds for tissue engineering. Technology and Health Care, 15(1), 3–17.

Odabaş, S., Sayar, F., Güven, G., Yanikkaya-Demirel, G., & Pişkin, E. (2008). Separation of mesenchymal stem cells with magnetic nanosorbents carrying CD105 and CD73 antibodies in flow-through and batch systems. Journal of Chromatography B, 861(1), 74–80.

Odabas, S. (2016). Collagen–carboxymethyl cellulose–tricalcium phosphate multi-lamellar cryogels for tissue engineering applications: Production and characterization. Journal of Bioactive and Compatible Polymers, 31(4), 411–422.

Rauh, J., Milan, F., Günther, K. P., & Stiehler, M. (2011). Bioreactor systems for bone tissue engineering. Tissue Engineering Part B: Reviews, 17(4), 263–280.

Sensenig, R., Sapir, Y., MacDonald, C., Cohen, S., & Polyak, B. (2012). Magnetic nanoparticle-based approaches to locally target therapy and enhance tissue regeneration in vivo. Nanomedicine, 7(9), 1425–1442.

Subramanian, M., Miaskowski, A., Jenkins, S. I., Lim, J., & Dobson, J. (2019). Remote manipulation of magnetic nanoparticles using magnetic field gradient to promote cancer cell death. Applied Physics A, 125(4), 1–10.

Vrana, N. E., Builles, N., Kocak, H., Gulay, P., Justin, V., Malbouyres, M., … & Hasirci, V. A. S. I. F. (2007). EDC/NHS cross-linked collagen foams as scaffolds for artificial corneal stroma. Journal of Biomaterials Science, Polymer Edition, 18(12), 1527–1545.

Vural, A. C., Odabas, S., Korkusuz, P., Yar Sağlam, A. S., Bilgiç, E., Çavuşoğlu, T., … & Vargel, İ. (2017). Cranial bone regeneration via BMP-2 encoding mesenchymal stem cells. Artificial cells, nanomedicine, and biotechnology, 45(3), 544–550.

Yeatts, A. B., & Fisher, J. P. (2011). Bone tissue engineering bioreactors: dynamic culture and the influence of shear stress. Bone, 48(2), 171–181.

Whitney, K. E., Liebowitz, A., Bolia, I. K., Chahla, J., Ravuri, S., Evans, T. A., … & Huard, J. (2017). Current perspectives on biological approaches for osteoarthritis. Annals of the New York Academy of Sciences, 1410(1), 26–43.

Williams, D. (2004). Benefit and risk in tissue engineering. Materials Today, 7(5), 24–29.

Zablotskii, V., Polyakova, T., Lunov, O., & Dejneka, A. (2016). How a high-gradient magnetic field could affect cell life. Scientific reports, 6(1), 1–13.

