## Supplemental Figure I for "Magnetically stimulated cryogels to enhance osteogenic and chondrogenic differentiation of stem cells"

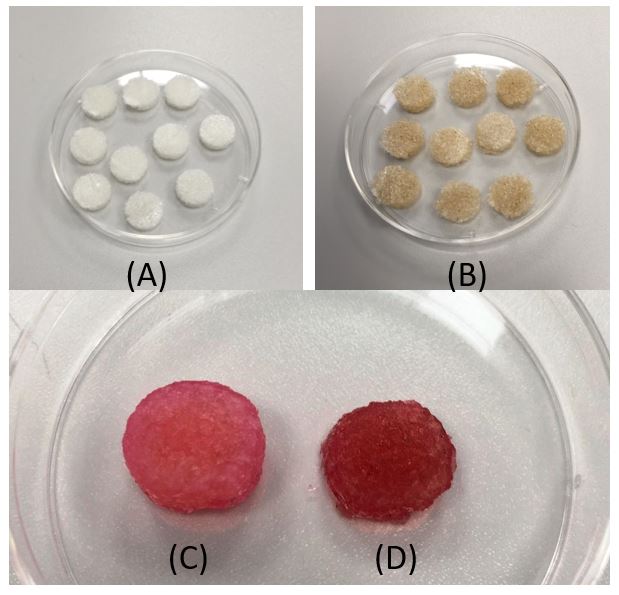

Supplementary Figure I. (A) Plain gelatin cryogels (B) Magnetic gelatin cryogels (C-D) Swollen Gel and M-Gel cryogels, respectively
